# Exploring the mechanism and pattern of bone formation during RANKL inhibition in a mouse model of fibrous dysplasia

**DOI:** 10.1101/2025.08.26.672317

**Authors:** G Farinacci, I Coletta, B Palmisano, E Spica, C Tavanti, S Di Cristofano, S Donsante, M Serafini, T Borsello, MA Venneri, A Corsi, D Salerno, B Donati, A Ciarrocchi, D Raimondo, M Riminucci

## Abstract

Fibrous dysplasia (FD) of bone is a genetic fibro-osseous disorder with increased bone remodeling activity. Inhibition of RANKL modifies FD lesions by inducing the replacement of the fibrous tissue with bone. This effect was observed in FD murine models receiving anti-mouse RANKL antibodies or small molecule RANKL inhibitors and in FD patients treated with denosumab. However, in neither case the mechanism and pattern of deposition of the newly formed bone were clarified. We performed radiographic, morphological and molecular analyses on EF1α-Gsα^R201C^ (FD) mice receiving an anti-mouse RANKL antibody. We observed that RANKL inhibition caused a decrease in the expression of genes involved in osteogenesis, osteoclastogenesis, matrix remodeling and osteoblast-osteoclast cross-talk in affected skeletal segments. Nonetheless, intra-lesional bone surfaces were covered by a continuous layer of osteoid, indicating that bone formation was actively ongoing in the pathological tissue in spite of the treatment. Accordingly, all bone surfaces within FD lesions showed calcein labeling which was never detected in the fibrous tissue far from bone. These results indicate that in the absence of RANKL activity, bone formation in FD tissue does not occur diffusely or stochastically. In contrast, it is restricted to bone surfaces where osteoprogenitor cells are orderly recruited from the adjacent fibrosis, progressively converting it into bone. Clinically, this suggests that the volume of pre-treatment bone in FD lesions may be a determinant of the skeletal improvement that individual patients may achieve during the same denosumab treatment course. As a consequence, it may also be a determinant of the severity of the rebound effect that they can experience upon treatment discontinuation.

## Introduction

Fibrous dysplasia of bone (FD, OMIM 174800) is a genetic skeletal disease due to mutations at *GNAS*, which encodes the α subunit of the stimulatory G protein (Gsα) ^1^. Most frequently, *GNAS* mutations cause the substitution of the arginine 201 in the Gsα protein (UNIPROT ID: P63092) with a cysteine (R201C) or a histidine (R201H), thereby reducing its GTPase function and leading to persistent activity of the cAMP signalling pathway. FD lesions result from the disrupted homeostasis of Gsα mutated osteoprogenitor cells in the post-natal bone marrow, which translates into the growth of a fibrotic stroma that produces hypo-mineralized bone while recruiting osteoclast precursors ^2^. The pathological fibro-osseous tissue replaces normal bone and marrow at one or multiple sites, according to the somatic mosaicism caused by the post-zygotic occurrence of the mutation, and results in mechanically incompetent skeletal segments that deform and fracture and often cause untreatable pain ^3–5^.

Although the deposition of pathological bone is the defining and best-known feature of FD ^6^, bone resorption typically outpaces bone formation within individual lesions, making FD a disease with locally unbalanced bone remodeling. Thus, a rational and widely used pharmacological treatment consists in the administration of anti-resorptive drugs. Among these, bisphosphonates have been shown to provide some relief from FD pain ^3^. However, neither in FD patients nor in mouse models, significant effect has been observed on the histopathology and progression of the disease ^7,8^. Conversely, both preclinical and clinical studies have shown that inhibition of Receptor activator of nuclear factor kappa-B ligand (RANKL), one of the most powerful osteoclastogenic cytokines ^9,10^ abundantly expressed by the FD tissue ^11–13^, reduces the bone turnover rate and the growth of lesions and, in some patients, also improves bone pain ^8,13–17^. Interestingly, despite the downregulation of bone remodeling, intra-lesional bone formation is not suppressed by RANKL inhibition. Indeed, in both transgenic mice ^8,13,14,17^, and FD patients, ^16,17^ lesions are variably refilled with hyper-mineralized bone, suggesting that the Gsα mutated stromal cells complete their differentiation into functional osteoblasts. However, the fine mechanisms leading to the replacement of the fibrous tissue and the pattern of new bone formation remain unclear.

In this study, we analysed FD lesions of EF1α-Gsα^R201C^ mice treated with different doses of an anti-mouse RANKL antibody. Our results demonstrate that terminal osteogenic differentiation of lesional progenitor cells during RANKL inhibition does not occur in a diffuse or stochastic manner and rather follows an ordered spatial pattern. Indeed, as shown by calcein labeling and von Kossa stain, *de novo* bone matrix deposition proceeds from the surface of pre-existing intra-lesional bone trabeculae to progressively replace the fibrous tissue. Translated into a clinical scenario, this pattern suggests that in FD patients, the density of bone trabeculae within each lesion may be an important determinant of the amount of bone therein produced during the treatment. Furthermore, our study shows that RANKL inhibition in mice significantly upregulates the expression of *Mmp2* in undifferentiated FD osteogenic cells, both *in vivo* and *in vitro*. Based on the involvement of this MMP in bone matrix mineralization, our result suggests a new potential molecular mechanism linking RANKL to bone matrix homeostasis.

## Materials and Methods

### Mice and experimental groups

The study was conducted in compliance with relevant Italian laws and Institutional guidelines and all procedures were approved by Institutional Animal Care and Use Committee (IACUC). EF1α-Gsα^R201C^ transgenic mice (FD mice), were generated as previously described ^18^ and genotyped using the oligonucleotide sequences reported in Table 1. All mice were maintained in cabin-type isolators at standard environmental conditions (temperature 22-25 °C) with 12:12 h dark/light photoperiod and food and water provided *ad libitum*. Three-month-old FD mice were treated with 300 µg/dose of anti-mouse RANKL antibody (αRANKL, clone IK22/5, Bio X Cell, West Lebanon, NH, USA) or vehicle (Veh, consisting in PBS) administered intraperitoneally (Fig. 1a). The treatment schedule included two doses in the first week (day 1 and 5), one dose in the second week (day 8) and another one in the third week (day 14). To monitor the intra-lesional morphological changes induced by αRANKL, mice were sacrificed either at day 3 (after 1 dose), at day 7 (after 2 doses), day 10 (after 3 doses) or day 17 (after 4 doses). Molecular studies were then performed on paraffin-embedded bone segments of mice sacrificed at day 17 (4 doses). To better highlight the sites of new bone deposition during RANKL inhibition, two of the five mice treated with 4 doses of αRANKL, also received two injections of 30 mg/kg calcein (Sigma-Aldrich, Saint Louis, MO, USA) in 2% sodium bicarbonate solution at 5 and 2 days before the sacrifice, which was carried out for all mice by carbon dioxide inhalation.

**Table 1.**
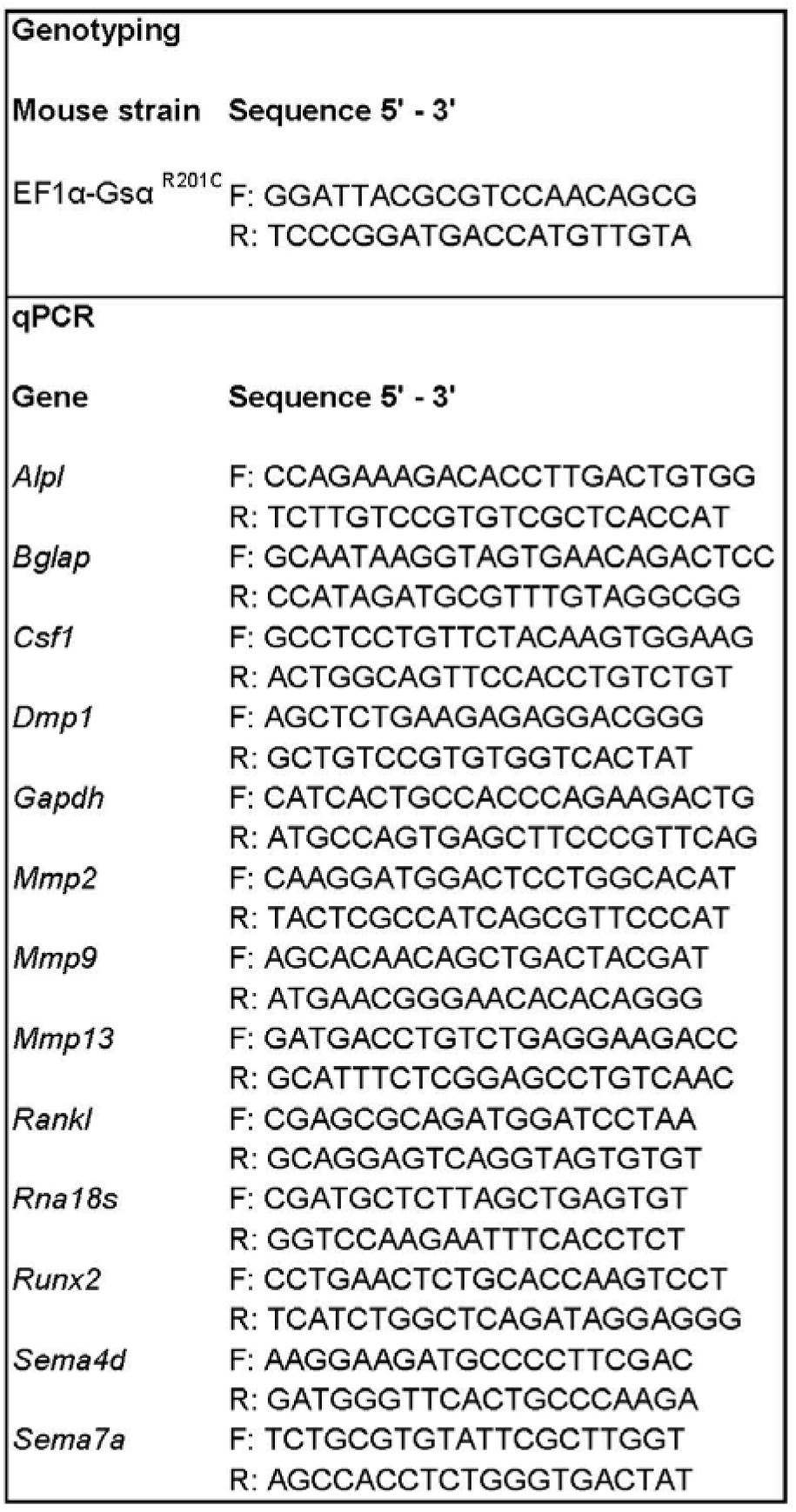
Sequence of primers used for genotyping and qPCR analysis.

**Fig. 1.**
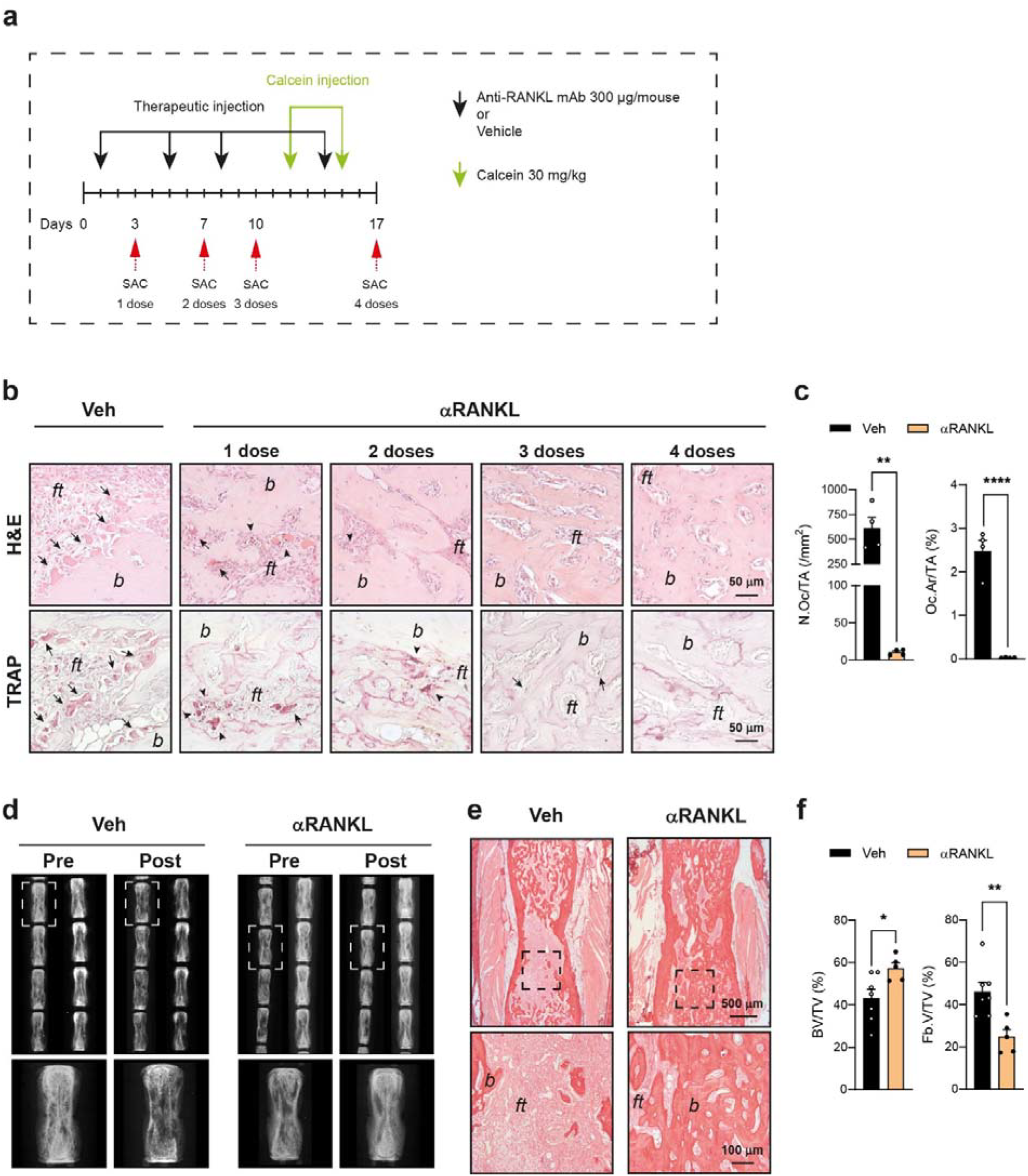
**a**) Experimental scheme of αRANKL or Vehicle (Veh) administration in EF1α-Gsα^R201C^ mice. **b**) Representative H&E and TRAP histochemistry on tail vertebrae from EF1α-Gsα^R201C^ mice receiving vehicle (Veh) or different doses of anti-RANKL antibody (αRANKL). Intra-lesion osteoclasts (arrows) become apoptotic (arrowheads) after 1 dose of αRANKL and progressively reduce after further doses. **c**) Quantification of osteoclast parameters showing a significant reduction after 4 αRANKL doses compared to Veh-treated EF1α-Gsα^R201C^ mice. N.Oc/TA: number of osteoclasts per tissue area; Oc.Ar/TA: osteoclast area per tissue area. Veh n=4; αRANKL n=4. Data are represented as the mean ± SEM. Statistical analysis was performed using Student t-test; ^**^ P < 0.01, ^****^ P < 0.0001. *b* = bone; *ft* = fibrous tissue. **d-f**) Replacement of FD tissue with bone in the tail vertebrae of mice receiving 4 αRANKL doses. Radiographic analysis demonstrates the increased density of affected vertebrae at the end (Post) compared to the beginning (Pre) of the treatment (**d**). Sirius red-stained tissue section (**e**) and histomorphometric analysis (**f**) confirm the higher amount of bone and lower amount of fibrous tissue in these samples compared to those from Veh-treated EF1α-Gsα^R201C^ mice. BV/TV: bone volume per tissue volume; Fb.V/TV: fibrous tissue volume per tissue volume. Veh n=7; αRANKL n=5. Data are represented as the mean ± SEM. Statistical analysis was performed using Student t-test; ^*^ P < 0.05, ^**^ P < 0.01. *b* = bone; *ft* = fibrous tissue.

### X-ray analysis

Radiographic analyses were performed at the beginning of the treatment and at the time of sacrifice using Faxitron MX-20 Specimen Radiography System (Faxitron X-ray Corp., Wheeling, IL, USA) set at 24 kV for 6 seconds, under anesthesia with a mixture of Zoletil 20 (Virbac SA, Carros, France) and Rompun (Bayer, Leverkusen, Germany).

### Histology and Histochemistry

Skeletal segments were fixed with 4% formaldehyde in PBS pH 7.4 for 48 hours at 4 °C and processed for either paraffin or methyl methacrylate (MMA) embedding.

For paraffin embedding, samples were decalcified with 10% EDTA and processed according to standard procedures. Three µm-thick sections were used for histological staining with Haematoxylin-Eosin (H&E), Sirius red, May Grünwald-Giemsa (MGG) and tartrate-resistant acid phosphatase (TRAP) histochemistry using Sigma Aldrich reagents (Sigma Aldrich) as previously described ^19^. Paraffin embedded tail vertebrae to be used for RNA extraction and NanoString analysis were selected based on histologic evaluation. MMA embedding was performed as described previously ^20^. Briefly, samples were fixed with 4% formaldehyde solution for 24 hours, dehydrated through increasing ethanol concentrations, infiltrated for 3 days with a plastic embedding solution containing MMA and then incubated at −20°C in a polymerization mixture obtained by adding N,N-dimethyl-p-toluidine to the embedding solution. Five-to-eight µm-thick sections were cut from each block and after removal of MMA with 2-methoxyethylacetate, stained with Toluidine Blue before calcein fluorescence evaluation or Von Kossa/Van Gieson to assess bone mineralization. All chemical reagents were purchased from Sigma Aldrich. Toluidine Blue stained sections were analyzed by an optical microscope (Zeiss Axiophot, Jena, Germany) and calcein-fluorescence was detected by a Leica Confocal Microscope (Wetzlar, Germany).

### Histomorphometry

Quantitative bone histomorphometry was conducted on tail vertebrae in a region of interest (ROI) between the two growth plates of tail vertebrae. The experiments were performed in a blinded fashion. Sirius red-stained sections were used to measure trabecular bone volume per tissue volume (BV/TV) and fibrous tissue volume per tissue volume (Fb.V/TV). TRAP-stained sections were used to measure osteoclast number per tissue area (N.Oc/TA) and osteoclast area per tissue area (Oc.Ar/TA). MGG-stained sections were used to measure osteoblast surface per bone surface (Ob.S /BS). Images were acquired with an optical microscope (Zeiss Axiophot, Jena, Germany) and all histomorphometric analyses were performed using ImageJ software (NIH, Bethesda, MD, USA). Results were reported using standard nomenclature and abbreviations ^21^.

### Immunohistochemistry

Immunolocalization of matrix metalloproteinase 2 (MMP2) and osterix (OSX) was carried out using rabbit anti-mouse antibodies (anti-MMP2 #GTX104577, GeneTex, Irvine, CA, USA; anti-OSX #ab22552, Abcam, Cambridge, UK) applied at a dilution of 1:300 and 1:200, respectively, in PBS and incubated overnight at 4 °C. For MMP2 immunostaining, a heat-mediated antigen retrieval step with citrate buffer (pH 6) at 98 °C for 30 min was performed. After the application of the primary antibody, sections were repeatedly washed with PBS and incubated for 30 min with biotin-conjugated polyclonal swine anti-rabbit IgG (#E0353, Dako, Agilent, Santa Clara, CA, USA, 1:500 in PBS) and then exposed for 30 min to peroxidase-conjugated streptavidin (#P0397, Dako, Agilent, Santa Clara, CA, USA; 1:1000 in PBS). The peroxidase reaction was developed using DAB substrate kit (SK-4105, Vector Laboratories, Burlingame, CA, USA).

### Gene expression analysis by Nanostring nCounter technology on formalin-fixed-decalcified-paraffin-embedded (FFDPE) bone samples

Five to ten tissue sections (20 µm thickness) were obtained from decalcified, paraffine embedded tail vertebrae (two vertebrae per mouse). Total RNA was extracted using RNeasy FFPE Kit (#73504, Qiagen, Hilden, Germany) according to the manufacturers’ instructions. RNA concentration and purity were assessed by NanoDrop 2000 spectrophotometer (Thermo Fisher Scientific, Waltham, USA). Only samples meeting quality thresholds (A260/A280 ≥ 1.7 and A260/A230 ≥ 1.8) were included for downstream analysis.

Gene expression profiling was conducted using NanoString nCounter platform with a custom designed CodeSet targeting 54 genes involved in osteoclastogenesis, matrix remodeling, osteoblast-osteoclast cross-talk and osteogenesis (Table S1). Raw gene counts were analyzed using nSolver Analysis Software 4.0 (NanoString Technologies, Seattle, WA, USA) as previously described ^22^. Samples were selected based on NanoString quality control metrics: percentage of fields of view (FOV) read (>75%), binding density (between 0.05 and 2.25), positive control linearity (>0.95) and positive control limit of detection (>2).

Raw counts were normalized using the geometric mean of the three housekeeping genes (*Actb, Cltc, Cnot10*) that showed an average count >150, after validation against positive and negative controls. Normalized gene counts were log_2_ transformed and used for downstream analyses.

### Clustering analysis and Silhouette plot

Gene expression data were visualized using heatmap, and hierarchical clustering analysis of both samples and genes was performed with “heatmap.2” function from the *gplots* package in R (RStudio). To determine the optimal number of clusters for sample classification, silhouette plot analysis was performed using *fviz_nbclust* function from the *factoextra* and *NbClust* packages.

### Cell cultures and osteogenic differentiation

Osteogenic cells were isolated from tail vertebrae of 5-month-old WT (n=3) and FD (n=4) mice. Briefly, after removal of skin and tendons, tail vertebrae were placed in HBSS (Gibco, Thermo Fisher Scientific) with 2 mg/ml collagenase type II (Gibco, Thermo Fisher Scientific) and incubated in a water bath for 15 minutes at 200 relative centrifugal force (rcf), to remove the outer layer of periosteum. Then, they were minced and further digested with 2 mg/ml collagenase type II solution in HBSS and were incubated in a water bath at 100 rcf for 1 hour. After digestion, to obtain a single cell suspension, samples were filtered through a 100 µm nylon mesh, centrifuged at 1300 rcf for 5 minutes, resuspended in 10 ml of complete culture medium consisting of αMEM (Sigma-Aldrich) supplemented with 20% FBS (Thermo Fisher Scientific), 1% L-glutamine (L-gln, Sigma-Aldrich) and 1% penicillin/streptomycin (P/S, Sigma-Aldrich) medium, filtered again through a 70 µm nylon mesh and centrifuged at 1300 rcf for 6 minutes. Cells were resuspended in complete culture medium and grown in a 100 mm culture dish. For osteogenic differentiation, cells were plated in 12-well plates at the density of 3 x 10^4^ cells/well and incubated for 14 days in medium containing complete αMEM supplemented with 4 mM β-glycerophosphate (Sigma-Aldrich) and 50 μg/ml of Ascorbic Acid (Sigma-Aldrich). Undifferentiated and differentiated cells were treated with either 1 μg/ml of αRANKL Ab or Veh for 6 hours in FBS-free medium.

### qPCR analysis

Quantitative PCR (qPCR) was performed on both fresh tissues from tail vertebrae and cells induced to osteogenic differentiation in vitro. Tail vertebrae were snap frozen in liquid nitrogen and homogenized by using Mikro-Dismembrator U (Gottingen, Germany). Total RNA was isolated from tissues and cells using the TRIzol Reagent protocol (Invitrogen, Thermo Fisher Scientific). Reverse transcription was performed by using Takara PrimeScript™ RT Reagent Kit with gDNA Eraser (Takara, Japan) according to the manufacturers’ instructions. qPCR analysis was conducted on cDNA samples by using a 7500 Fast Real-Time PCR System (Applied Biosystem, Waltham, Massachusetts, USA), performed using PowerUP Sybr Green (Thermo Fisher Scientific) and gene expression levels of each gene were normalized to *Gapdh* or *Rna18s* expression. The primers used are listed in Table 1.

### Statistical analysis

One-way ANOVA followed by a Tukey’s multiple comparison test was used to compare three groups while the comparisons between two groups were performed using the unpaired Student t-test. In all experiments a P-value less than 0.05 was considered statistically significant. All graphs and statistical analyses were performed using GraphPad Prism version 8 (GraphPad Software, La Jolla, CA, USA).

## Results

### Intra-lesion bone deposition follows osteoclast reduction in RANKL inhibited FD mice

Three-month-old FD mice with radiographic evidence of skeletal lesions were sacrificed after receiving different doses of αRANKL or Veh (Fig. 1a). Tail vertebrae were harvested and analyzed in order to assess the morphological effects of RANKL inhibition on the FD tissue and to identify lesions with ongoing fibrous to bone replacement. The earliest tissue changes induced by αRANKL treatment consisted in the appearance of apoptotic bodies in both mono and multi-nucleated TRAP positive cells, as observed in the vertebrae of mice that received either one or two doses (Fig.1b). The number of multinucleated osteoclasts started to decline after the third αRANKL dose and was significantly reduced compared to the Veh-group in FD mice receiving the four-dose treatment (Fig. 1b,c) when the process of new bone formation became evident. Indeed, comparison of radiographic images of individual mice taken at the beginning and at the end of αRANKL administration showed a reduction in the number and size of lytic areas (Fig.1d) whereas histomorphometry revealed significant differences in the amount of bone (increased) and fibrous tissue (decreased) in lesions of αRANKL-compared to Veh-treated FD mice (Fig.1e,f). Based on these findings, tail vertebrae from the four-dose experimental group were selected for subsequent NanoString gene expression profiling.

### RANKL inhibition downregulates osteogenic genes and increases Mmp2 in mouse FD lesions

Following the identification of αRANKL vertebrae with ongoing refilling of lytic areas, we analyzed the corresponding samples to evaluate treatment-induced changes in gene expression. To this aim, total mRNA was extracted from FFDPE tail vertebrae of αRANKL- and Veh-treated FD mice and untreated wild-type (WT) littermates and analyzed using NanoString nCounter technology with a multiplex Custom CodeSet. The selected genes were relevant to osteoclastogenesis, matrix remodeling, osteoblast-osteoclast cross-talk and osteogenesis.

Unsupervised clustering (Fig. 2a) and silhouette plot analyses (Fig. S1a) revealed two distinct clusters, one corresponding to Veh-treated FD mice and the other including αRANKL-treated FD and WT mice. Compared to WT, Veh-treated FD mice exhibited increased expression of several bone remodeling-related genes, including *Csf1, Rankl, Ctsk* and *Sema4d*, while other transcripts such as *Sema7a, C3* and *Tln1* remained unchanged (Fig. 2b). Furthermore, *Mmp9* and *Mmp13* mRNAs were also upregulated in Veh-treated FD samples, whereas *Mmp2* mRNA remained unchanged despite lesion development. Osteogenic differentiation markers were also elevated, with the exception of *Bmp6* and *Atf4*, which were decreased compared to WT (Fig. 2c). Expression of the two adipogenic markers, *Pparg* and *Fabp4*, was markedly lower than WT, consistent with the absence of marrow adipocytes within the FD tissue (Fig. 2c).

**Fig. 2.**
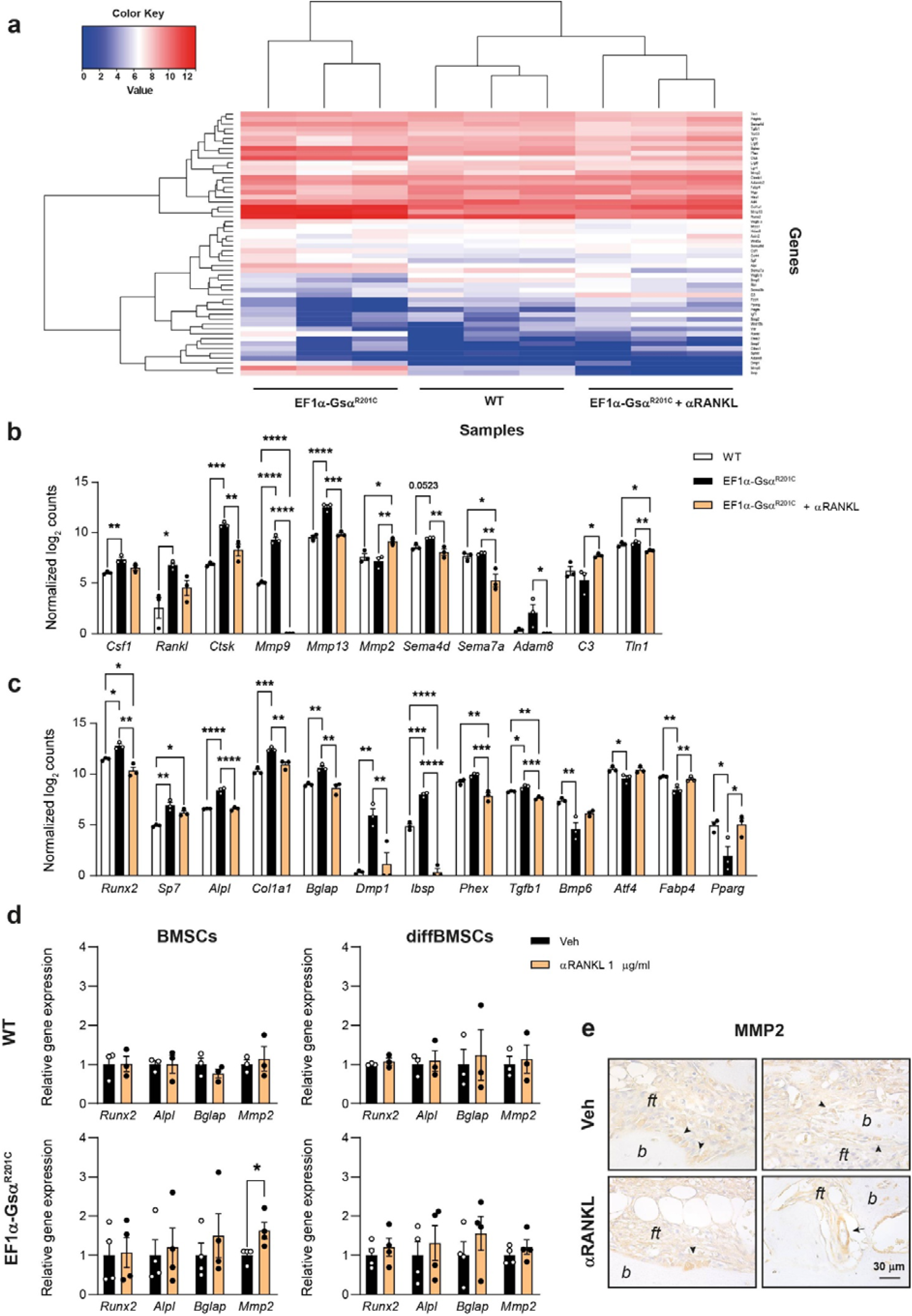
**a**) Heatmap representation of gene expression data obtained from FFDPE tail vertebrae. Clustering analysis showing two clusters, one corresponding to Veh-treated EF1α-Gsα^R201C^ mice and the other one corresponding to WT and αRANKL-treated EF1α-Gsα^R201C^ mice. The relative abundance of transcripts is indicated by the colour (blue, lower; red, higher). **b**,**c**) Column bars representation of gene expression analysis of genes related to osteoclasts, matrix remodeling, osteoblast-osteoclast crosstalk (**b**) and genes involved in osteogenic and adipogenic differentiation (**c**) in WT and EF1α-Gsα^R201C^ mice treated with Veh or αRANKL. WT n=3; EF1α-Gsα^R201C^ n=3; EF1α-Gsα^R201C^ + αRANKL n=3. Data are represented as the mean ± SEM. Statistical analysis was performed using One-Way ANOVA followed by a Tukey’s multiple comparison test; ^*^ P < 0.05, ^**^ P < 0.01, ^***^ P < 0.001, ^****^ P < 0.0001. The exact P-value was reported on *Sema4d* column bars. **d**) qPCR performed on undifferentiated (BMSCs) and differentiated (diffBMSCs) stromal cells isolated from tail vertebrae of WT and EF1α-Gsα^R201C^ mice and treated with Veh or 1 µg/ml αRANKL. A significant change was observed only in the level of expression of *Mmp2* in EF1α-Gsα^R201C^ BMSCs. WT: Veh n=3, αRANKL 1 µg/ml n=3; EF1α-Gsα^R201C^: Veh n=4, αRANKL 1 µg/ml n=4. Data are represented as the mean ± SEM. Statistical analysis was performed using Student t-test; ^*^ P < 0.05. **e**) Immunolocalization of MMP2 in EF1α-Gsα^R201C^ mice treated with Veh or αRANKL showing the expression of MMP2 in some osteoblasts (arrowheads) bordering the bone surface in both Veh- and αRANKL-treated mice and in perivascular stromal cells (arrows) only in mice treated with αRANKL. *b* = bone; *ft* = fibrous tissue.

Following αRANKL treatment, osteoclast-derived gene transcripts became low or undetectable, although *Csf1* and *Rankl* expression was not affected (Fig. 2b). Among matrix remodeling genes, *Mmp9* expression was nearly abolished and *Mmp13* returned to the WT levels. Interestingly, *Mmp2*, a metalloproteinase implicated in bone mineralization and density ^23,24^, was significantly upregulated in αRANKL-treated FD vertebrae compared with both Veh-treated FD and WT mice (Fig. 2b). Despite histological evidence of intra-lesion bone deposition, osteogenic genes expression was reduced in αRANKL-compared to Veh-treated FD mice, and in some cases even lower than WT samples. Of note, this reduction involved both early (*Runx2* and *Alpl*) and late (*Dmp1* and *Phex*) osteogenic transcripts (Fig. 2c). Furthermore, in parallel with the decrease in the amounts of osteogenic transcripts, the αRANKL treatment restored the levels of mRNAs for *Pparg* and *Fabp4* (Fig. 2c), in agreement with the focal evidence of developing multilocular adipocytes in this group of vertebrae (Fig. S2a)

NanoString results were validated by qPCR analysis of selected genes using fresh vertebral tissue from 3-month-old FD and WT mice (Fig. S1b).

### αRANKL stimulates Mmp2 expression in mouse FD osteoprogenitor cells

To assess whether αRANKL directly modulated gene expression in FD cells, we performed *in vitro* studies. Cells isolated from the tail vertebrae of Veh-treated FD mice and WT littermates were incubated in either standard growth medium or osteogenic medium for 14 days and treated with 1 µg/ml of αRANKL for 6 hours. qPCR analysis was then carried out to assess the level of expression of representative osteogenic genes, including *Runx2* and *Alpl* for early osteogenic commitment and *Bglap* for late stages of osteogenic differentiation. Furthermore, we analyzed the expression of *Mmp2*. Across all conditions αRANKL treatment had no significant impact on the expression of the osteogenic transcripts (Fig. 2d). However, these experiments showed that αRANKL induced a significant increase in *Mmp2* mRNA in undifferentiated FD stromal cells, although no effect was observed after osteoblast-like differentiation (Fig. 2d). In agreement with this finding, immunolocalization of MMP2 showed a positive signal in some undifferentiated perivascular stromal cells only in bone lesions of αRANKL-treated FD mice, whereas MMP2 expression in osteoblasts was overall comparable between αRANKL- and Veh-treated FD mice (Fig. 2e).

### Bone formation in mouse FD lesions during RANKL inhibition is restricted to bone surfaces

*In vivo* and *in vitro* molecular studies suggested that inhibition of RANKL in FD mice did not associate with a massive osteogenic differentiation of cells within the intra-lesional fibrous tissue. This conclusion was further supported by histomorphometry and immunophenotypic evaluation performed on the same FFDPE vertebrae used for gene expression analysis. Indeed, MGG stain did not show changes in the morphology and/or organization of cells within the fibrotic tissue (Fig.3a). In contrast, osteoblast-like cells remained restricted to bone surfaces with no histomorphometric differences compared to Veh-treated FD vertebrae (Fig. 3a,b). Furthermore, immunolocalization experiments confirmed that, as in Veh-treated FD samples, the expression of the transcription factor OSX remained localized to the osteogenic cells bordering the bone trabeculae (arrowheads; Fig. 3c), with only a few, sparse positive cells within the fibrous tissue (arrows; Fig. 3c). Based on these observations, calcein labeling was performed to further clarify the pattern of bone matrix deposition. In the tail vertebrae (Fig. 4a) and femurs (Fig. 5a) of the Veh group, the green fluorescence was bright, extensive and smeared on all lesional bone trabeculae. This appearance was consistent with the florid deposition of bone, related to the enhanced bone remodeling, with osteomalacic changes that typically occur in FD, both in mice ^18^ and patients ^25,26^.

**Fig. 3.**
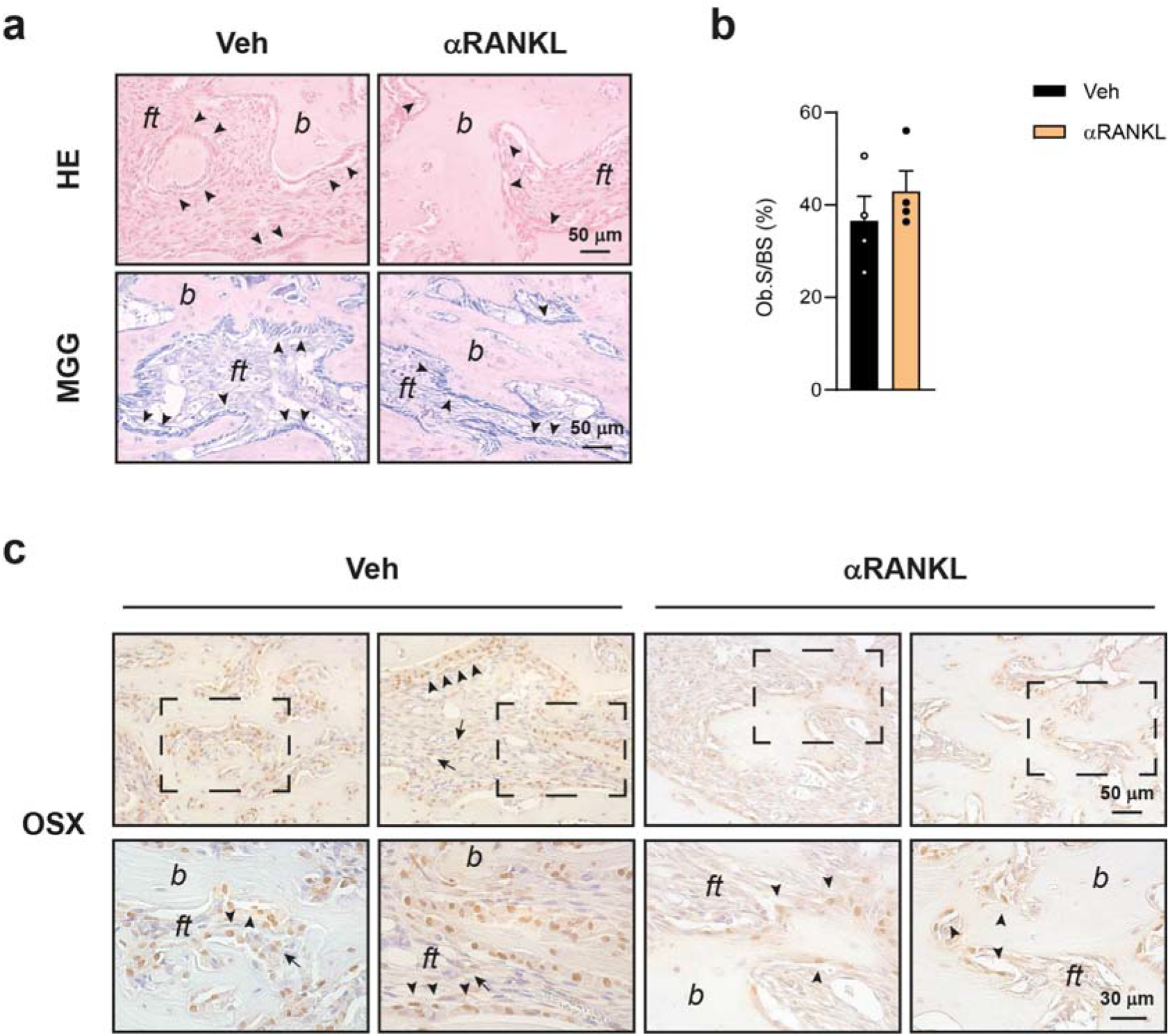
**a**) H&E and MGG stains showing osteoblasts (arrowheads) within tail lesions of αRANKL-treated EF1α-Gsα^R201C^ mice. **b**) Histomorphometric analysis reveals comparable values of osteoblast surface per bone surface (Ob/S/BS) in the tail vertebrae of αRANKL-compared to Veh-treated EF1α-Gsα^R201C^ mice. Veh n=4; αRANKL n=4. Data are represented as the mean ± SEM. Statistical analysis was performed using Student t-test. **c**) Immunolocalization of OSX in EF1α-Gsα^R201C^ mice treated with Veh or αRANKL showing the expression of OSX in osteoblasts (arrowheads) bordering the bone surface and in a few cells within fibrous tissue (arrows). *b* = bone; *ft* = fibrous tissue.

**Fig. 4.**
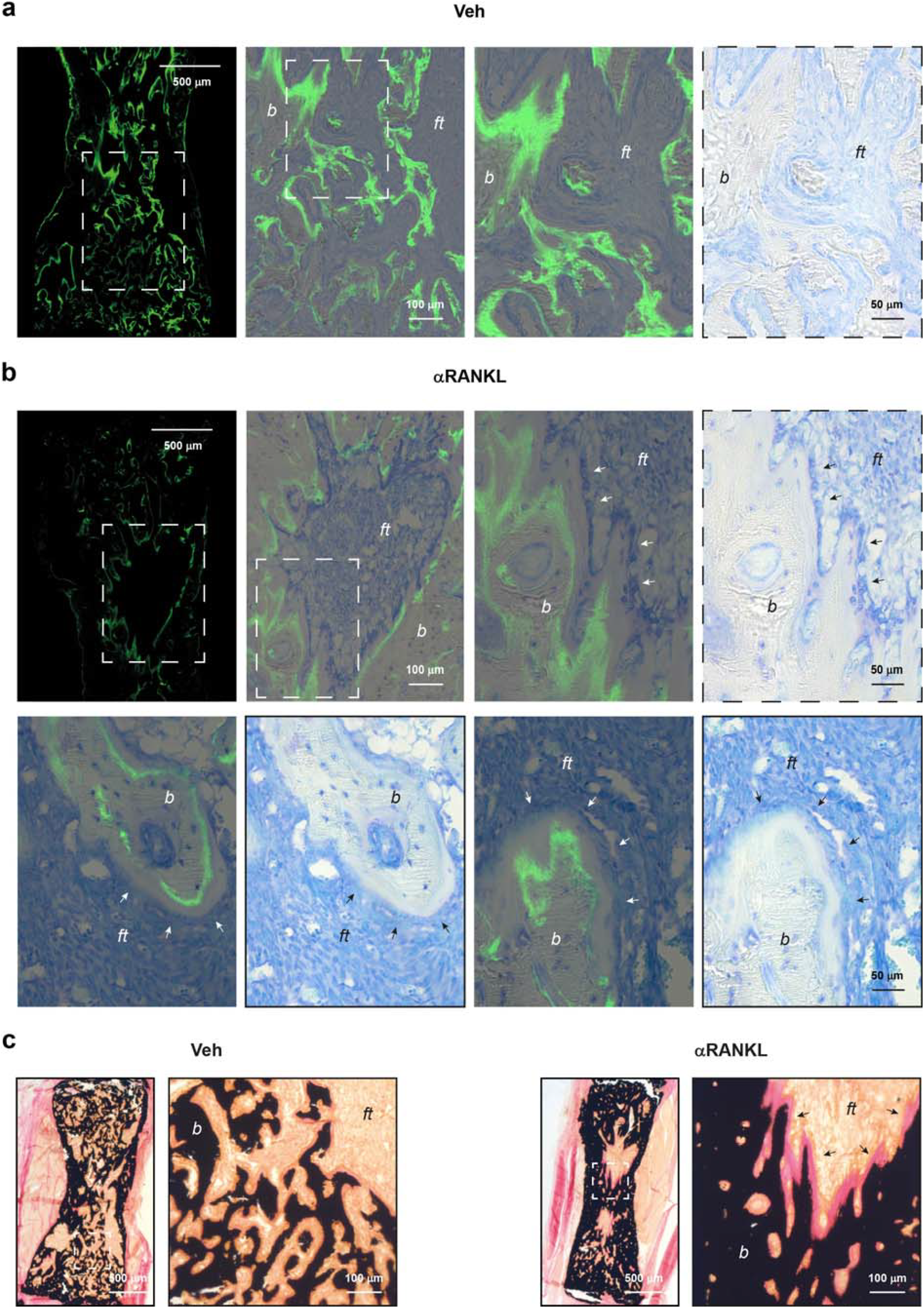
**a-b**) Tail vertebrae of EF1α-Gsα^R201C^ mice labeled with calcein and stained with Toluidine Blue. Fluorescence microscopy reveals a smeared pattern of fluorescence in Veh-treated EF1α-Gsα^R201C^ mice (**a**). After αRANKL treatment, calcein labeling is detected as a single line, with focally smeared areas, on all intra-lesional bone surfaces where newly formed bone bordered by osteoblasts (arrows) is also observed (**b**). Note the absence of fluorescent labeling within the fibrous tissue. **c**) Von Kossa/Van Gieson-stained sections of undecalcified tail vertebrae showing the presence of osteoid seams (arrows) on the bone surface of αRANKL-treated EF1α-Gsα^R201C^ mice. *b* = bone; *ft* = fibrous tissue.

**Fig. 5.**
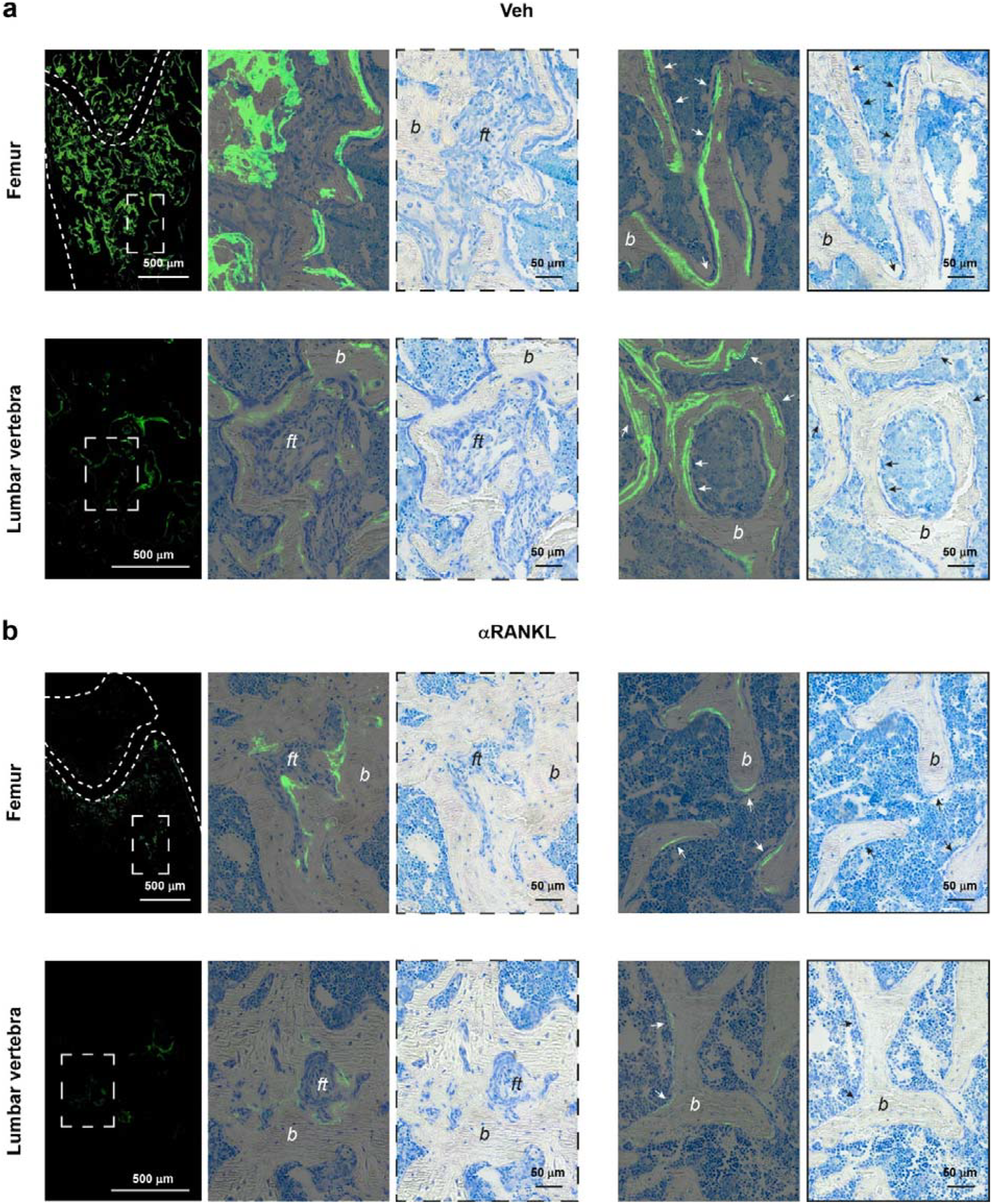
**a-b**) Femurs and lumbar vertebrae of EF1α-Gsα^R201C^ mice labeled with calcein and stained with Toluidine Blue. Numerous bright double fluorescent lines are observed on unaffected bone trabeculae of Veh-treated EF1α-Gsα^R201C^ mice (**a**, arrows). Calcein labelling is markedly reduced in αRANKL-treated EF1α-Gsα^R201C^ mice in which double fluorescent lines are rarely observed on unaffected bone trabeculae (**b**, arrows). *b* = bone; *ft* = fibrous tissue.

As expected, double labeling was observed on femoral bone trabeculae spared from the disease and in unaffected lumbar vertebrae (Fig. 5a). In all the examined skeletal segments, the intensity and distribution of the fluorescent labeling was overall reduced in αRANKL-treated compared to Veh-treated mice (Fig. 4b and Fig. 5b), reflecting the downregulation of bone remodeling and the consequent reduction of resorption surfaces available for refilling by new bone. However, within the pathological tissue, calcein staining was detected on all bone trabeculae, where it occurred as single fluorescent lines with focally smeared areas (Fig. 4b and Fig. 5b). In the same samples, Toluidine blue (Fig. 4b) and von Kossa stains (Fig. 4c) revealed the presence of osteoid seams covering the entire interface between the mineralized bone and the fibrous tissue. Of note, no deposit of the fluorochrome was ever observed within the fibrous tissue far from bone (Fig 4b and Fig. 5b). As in Veh-treated FD mice, unaffected bone trabeculae of femurs and lumbar vertebrae of αRANKL-treated FD mice showed double labeling, but the frequency of the staining was very low (Fig. 5b).

Overall, these results suggested that in αRANKL-treated FD mice, following refilling of resorption spaces, bone deposition continued on the bone surface facing the fibrous tissue and progressively replaced it. However, part of the bone matrix was still unmineralized at the end of the experiment, consistent with the presence of a single line of calcein labeling. The ordered occurrence of bone formation at the interface between bone surfaces and marrow fibrosis, was further supported by the evidence within lytic lesions, of a thick and continuous layer of bone matrix localized between the reversal lines, well highlighted by TRAP ^27^, and the remaining fibrous core (Fig. S2b).

Since the process of new bone formation was limited to the trabecular surfaces in contact with the fibrous tissue whereas activation of new remodeling sites was inhibited on all bone surfaces within the skeletal segment, altogether these results not only clarified the mechanisms of intra-lesional bone mass gain induced by RANKL inhibition but also explain why it associated with a reduction rather than an increase in the level of osteogenic transcripts and calcein labeling.

## Discussion

Inhibition of RANKL activity is currently the only therapeutic approach able to modify the tissue pathology of FD by inducing the replacement of fibrous tissue with bone. This effect has been observed in transgenic mouse models ^8,13,14,17^ and FD patients ^16,17^, and is induced by both antibodies ^8,13,16,17^ and small molecules that inhibit RANKL activity. ^14^ Notably, this effect is not replicated by other anti-bone resorption drugs that target osteoclast function rather than formation, such as BPs ^8^. However, few data are currently available on the mechanism of formation and pattern of deposition of the replacing bone matrix. In most clinical studies, FD patients’ response to the humanized anti-RANKL antibody denosumab, was assessed through surrogates for bone biopsy, such as serum levels of bone turn-over markers (BTMs) and bone imaging ^28–30^. However, serum BTMs, being evaluated on a systemic scale, reflect the effect of the drug on affected and non-affected skeletal segments rather than the osteogenic activity within individual FD lesions. Bone imaging, on the other side, captures treatment-dependent modifications in the density or tracer uptake of affected bones but does not reveal how the changes developed. Furthermore, patient monitoring usually starts at relatively long intervals from the beginning of the treatment, leaving its early effects unexplored. Similarly, in most of available studies on FD transgenic mice, histological evaluation was performed at the end of the treatment rather than during drug administration. A single, recent work reported histological and immunophenotypic evaluation of bone biopsies obtained from FD patients and bone samples from transgenic mice after 6 months and 4 weeks respectively of RANKL inhibition. In both cases, an increase in the number of SOST-positive osteocytes was observed, consistent with the increased amount of intra-lesional bone, whereas a reduction of *Runx2* and *Alpl* was observed in the mice ^17^.

In our study, we monitored FD lesions of EF1α-Gsα^R201C^ mice after different doses of an anti-mouse RANKL antibody. We first observed that a significant replacement of fibrous tissue with bone could be detected only after apoptosis of pre-existing TRAP positive osteoclasts that started shortly after the first dose and caused their significant reduction. Since it was previously reported that osteoclasts support the proliferation of FD osteoprogenitor cells ^17^ whereas osteoclast-derived apoptotic bodies promote osteogenesis ^31^, we hypothesized that the apoptotic removal of ectopic osteoclasts abundantly distributed within FD lesions ^32^ could lead to a diffuse process of terminal cell differentiation and bone deposition within the osteogenic fibrous tissue. However, the downregulation of osteogenic markers, which was in agreement with the results of previous study ^17^, and the absence of calcein labeling in the fibrotic areas far from bone surfaces, could be hardly reconciled with this possibility. Indeed, Von Kossa stain and calcein labeling demonstrated that during the anti-RANKL treatment, the osteogenic activity remained restricted to the bone surface. Indeed, refilling of pre-existing remodeling spaces can be considered as the first mechanism for the bone mass gain caused by the treatment. The refilling process is the main mechanism that underlies the effect of RANKL inhibition in denosumab-treated osteoporotic patients, in which osteogenic cells already engaged in bone formation continue to form extracellular matrix and complete the refilling of individual bone multicellular units (BMUs) before becoming quiescent ^33,34^. FD is a high bone remodeling disease in which the continuous activation of resorption sites shortens the life-span of pre-existing BMUs. In this context, the downregulation of bone remodeling caused by RANKL inhibition, confirmed by both the suppression of osteoclastogenic and osteogenic markers and the overall reduced calcein labeling, is expected to allow the refilling of an abnormally high number of BMUs, leading to a significant increase in bone mass that is not observed in osteoporosis. In addition to the BMU refilling process, our results suggest a second mechanism leading to bone matrix apposition. Specifically, they suggest a progressive recruitment and differentiation of osteogenic cells from the adjacent osteogenic fibrous tissue to the bone surface, a mechanism that would easily lead to the replacement of fibrous tissue with bone. Different hypotheses can be made to explain this process. For example, it is reasonable to assume that, compared to basal conditions, in RANKL inhibited FD lesions refilling of pre-existent resorption surfaces and lack of activation of new remodeling sites both increase the osteoconductive bone surface on which adjacent osteoprogenitor cells may convert into osteoblasts. In addition, it is possible that, although RANKL inhibition seems to reduce the overall levels of the mutated Gsα mRNA ^17^, the residual expression of the mutation in differentiated osteogenic cells may act as a further stimulator of bone formation, as reported in previous work ^35^. Regardless of the mechanisms, these results may help to better understand the factors influencing intra-lesion bone mass gain in FD patients treated with denosumab as well as the rebound phenomenon at treatment discontinuation, for which skeletal improvement is a major determinant along with the pre-treatment level of BTMs ^29^.

We previously showed in the same EF1α-Gsα^R201C^ transgenic mouse models, that the bone tissue produced during long-term RANKL inhibition is hyper-mineralized ^13^. In this study, part of the newly formed bone matrix produced within lesions was still unmineralized at the end of the experiment, as shown by the presence of a single calcein label in contrast with the double labeling of the unaffected bone. This suggests that the mineralization status achieved in the previous long-term RANKL inhibition study ^13^, likely depended on the long secondary mineralization phase due to the continuous inhibition of bone resorption ^33,34,36^. The factors controlling bone matrix mineralization during RANKL inhibition need to be better investigated in further studies. Meanwhile, our results suggest that MMP2, a matrix remodeling enzyme that is known to affect bone mineralization, ^23,24^ might be a potential candidate. Since MMP2 is normally produced by osteocytes and bone marrow stromal cells ^37^, the enhanced amount of its transcript in treated mice likely reflected the increased number of osteocytes, resulting from new bone formation, as well as the increased production by stromal osteoprogenitor cells upon stimulation by the anti-RANKL antibody, as revealed by *in vitro* study.

Our work provides additional findings relevant to the biology and treatment of FD. We observed that transcripts for *Rankl* and *Csf1*, which were significantly enriched in untreated FD mice ^17,38,39^, did not decrease during RANKL inhibition. This finding must be taken into account to better understand the rebound effect that may follow treatment withdrawn. Indeed, it is expected that during the administration of the anti-RANKL antibody, CSF1 continues to act on the monocyte lineage expanding the compartment of early committed osteoclastic cells and that a high number of mature osteoclasts is then rapidly generated from these cells when the anti-RANKL antibody is removed.

Finally, as expected, the adipogenic markers *Pparg* and *Fabp4* were downregulated in untreated FD lesions compared to WT bone. Following inhibition of osteoclastogenesis, their transcripts increased consistently with the focal histological evidence of developing adipocytes and in agreement with previous findings showing that adipose marrow was increased in anti-RANKL-treated FD mice compared to IgG-treated mice ^13^. The absence of adipocytes within FD lesions is commonly ascribed to the inability of Gsα mutated skeletal progenitors to undertake the adipogenic program. However, it was previously shown that in normal human progenitor cells transduced with the mutated Gsα sequence, adipogenic mRNA were significantly increased compared to control cells ^11^. Based on these published data and our current results, it may be hypothesized that the disappearance of adipose marrow within FD lesions may be due not only to the dysregulation of the intrinsic mechanisms regulating cell fate but also to the massive engagement of skeletal progenitor in osteogenesis at the expense of adipogenesis.

In conclusion, our study shows that during RANKL inhibition, new bone formation within FD lesions does not occur diffusely or stochastically within the fibrous tissue but follows an ordered pattern originating from bone surfaces, progressively replacing adjacent fibrous tissue. Clinically, these findings suggest that the amount of bone rather than fibrous tissue within FD lesions is a critical determinant of the bone mass gain that a given denosumab course may induce in FD patients and, as a consequence, of the severity of the rebound effect that may follow treatment discontinuation.

## Supporting information

Graphical Abstract

## Acknowledgements

Orphan Disease Center University of Pennsylvania in partnership with Fibrous Dysplasia Foundation (MDBR-19-110-FD, MDBR-21-110-FD, MDBR-23-010-FDMAS) to M.R.; Sapienza University (RP124190960A2E07 to AC).

## Data availability

Data will be available from the corresponding author upon request.

## Authorship

G.F. Conceptualization, Methodology, Data analysis, Writing original draft; I.C Methodology, Data analysis; B.P. Methodology, Data analysis; E.S. Methodology; C.T. Methodology; S.D.C. Data analysis; S.D. Methodology; M.S. Editing final version; T.B. Editing final version; M.A.V Editing final version; A.C. Editing final version; D.S. Data analysis; B.D. Data analysis; A.C. Data analysis; D.R. Conceptualization, Data analysis, Supervision; M.R. Conceptualization, Writing and Editing final version, Supervision.

All authors revised the paper critically for intellectual content, approved the final version and agreed to be accountable for the work.

