## Supplemental Figures for "Exploring the mechanism and pattern of bone formation during RANKL inhibition in a mouse model of fibrous dysplasia"

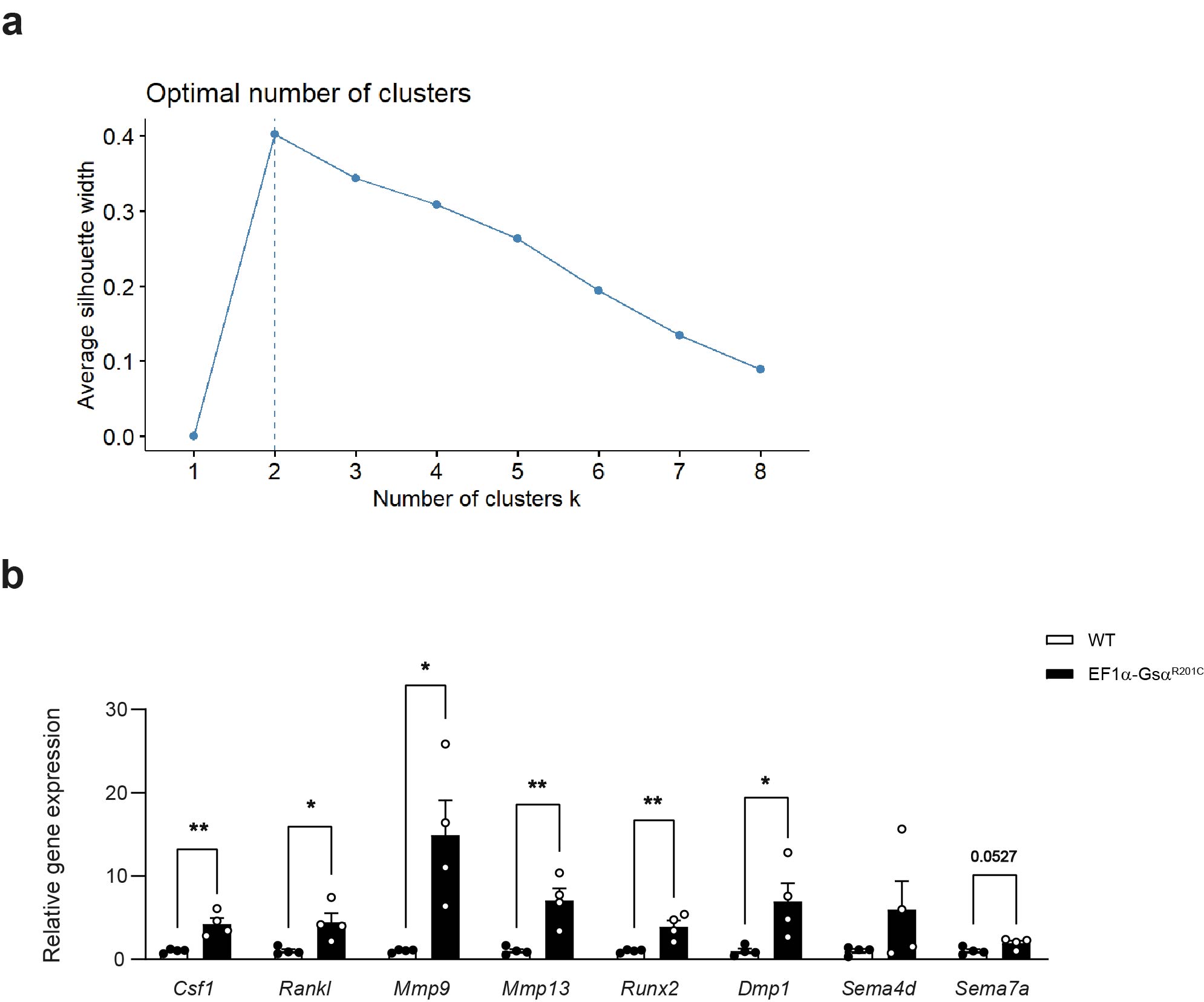

**Fig.S1** **a**) Silhouette plot analysis showing samples segregation into two clusters. **b**) Gene expression analysis performed on whole tail vertebrae of 3-month-old WT and EF1α-Gsα^R201C^ mice. WT n=4; EF1α-Gsα^R201C^ n=4. Data are represented as the mean ± SEM. Statistical analysis was performed using Student t-test; * P < 0.05, ** P < 0.01. The exact P-value was reported on *Sema7a* column bars.

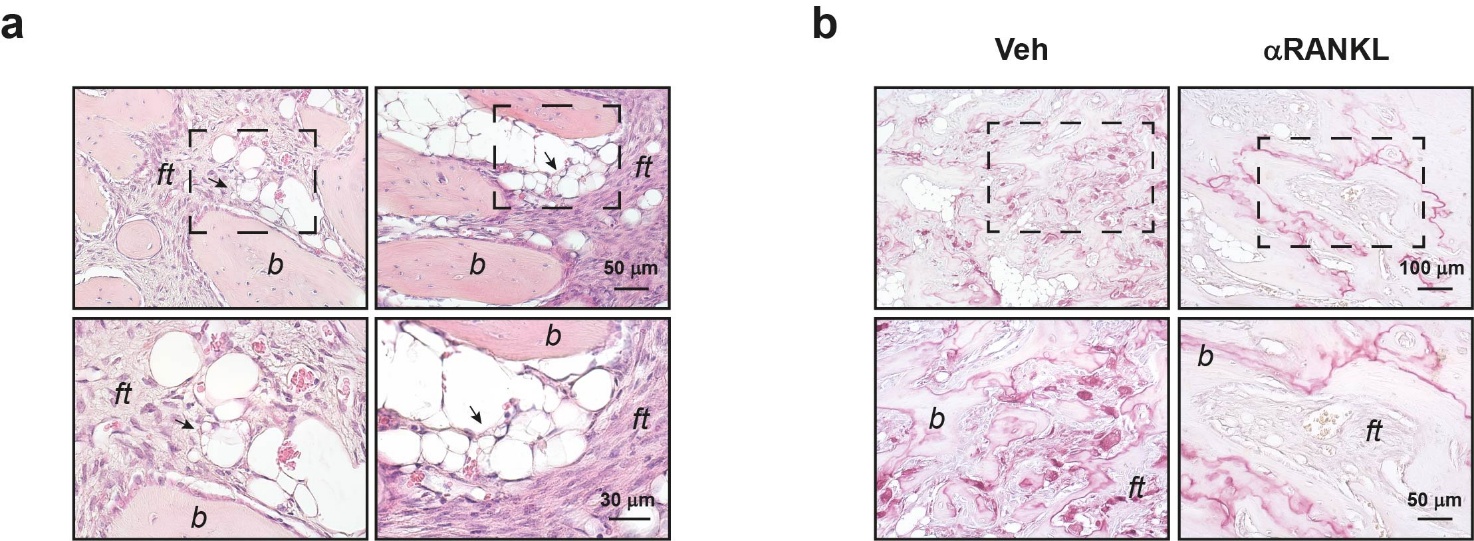

**Fig.S2** **a**) H&E-stained tissue sections of tail vertebrae of αRANKL-treated EF1α-Gsα^R201C^ mice showing the development of multilocular adipocytes (arrows). *b* = bone; *ft* = fibrous tissue. **b**) TRAP histochemistry of tail vertebrae showing the presence of multiple sites of bone remodeling in Veh- treated EF1α-Gsα^R201C^ mice and their reduction during αRANKL treatment. Note the bone produced on the TRAP positive reversal lines during RANKL inhibition encasing the residual fibrous tissue. *b* = bone; *ft* = fibrous tissue.
