## Supplementary figures and images for "Exploring the mechanism and pattern of bone formation during RANKL inhibition in a mouse model of fibrous dysplasia"

### Graphical Abstract

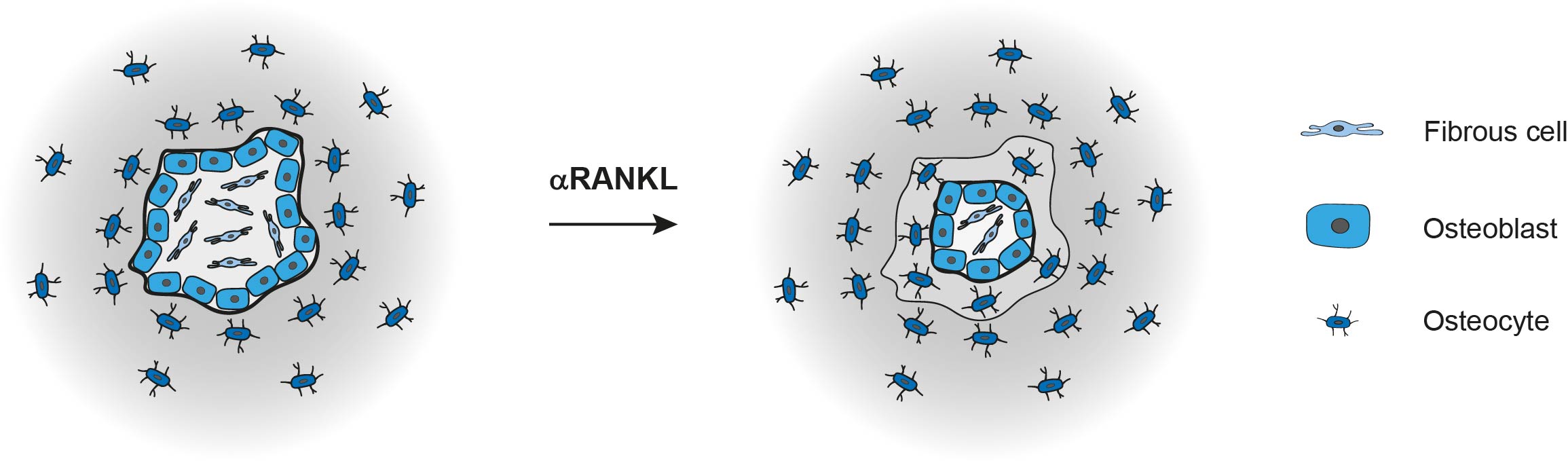
